# Glucose appetition in C57BL/6J mice: Influence of nonnutritive sweetener experience, food deprivation state and sex differences

**DOI:** 10.1101/2024.02.27.582331

**Authors:** Anthony Sclafani, Karen Ackroff

## Abstract

In addition to its sweet taste, glucose has potent and rapid postoral actions (appetition) that enhance its reward value. This has been demonstrated by the experience-induced preference for glucose over initially preferred nonnutritive sweetener solutions in 24-h choice tests. However, some sweetener solutions (e.g., 0.8% sucralose) have inhibitory postoral actions that may exaggerate glucose appetition whereas others (e.g., 0.1% sucralose + 0.1% saccharin, S+S) do not. Experiment 1 revealed that food-restricted (FR) male C57BL/6J mice displayed similar rapid glucose appetition effects (stimulation of glucose licking within minutes) and conditioned flavor preferences following 1-h experience with flavored 0.8% sucralose or 0.1% S+S and 8% glucose solutions. Thus, the inhibitory effects of 0.8% sucralose observed in 24-h tests were not apparent in 1-h tests. Experiment 2 evaluated the effects of food deprivation state on 1-h glucose appetition. Unlike FR female mice, ad libitum (AL) fed mice displayed no or delayed stimulation of glucose licking depending upon the training solutions used (0.1% S+S vs. 8% glucose, or 0.2% S+S vs. 16% glucose). Both AL groups, like the FR group, developed a preference for the glucose-paired flavor over the S+S paired flavor. Thus, food restriction promotes glucose appetition but is not required for a conditioned preference. Overall, male and female mice showed similar glucose appetition responses although females displayed a more rapid initial glucose response.

## 1. Introduction

Sugar is a very attractive nutrient that motivates eating and conditions preferences for associated flavors. Sugar appetite is mediated in part by the activation of oral sweet taste receptors (T1R2+T1R3 in mammals) and also by sugar-specific sensors in the intestinal tract [4,10]. The postoral appetite stimulating action of sugar is referred to as “appetition” to differentiate it from satiation effects which suppress appetite [13]. Sugar appetition was demonstrated in studies in which rodents were trained to drink a flavored nonnutritive solution (the CS+, e.g., grape) paired with a concurrent intragastric (IG) infusion of a sugar solution. Compared to an alternative flavored solution (the CS-, e.g., cherry) paired with IG water infusions, rats and mice showed increased intake of the CS+ in one-bottle sessions and strongly preferred the CS+ to the CS-in a two-bottle choice test [4]. This appetition response is sugar specific. In C57BL/6J (B6) mice, IG infusions of glucose and sucrose are very effective while fructose infusions are ineffective [14,22].

B6 mice also displayed quite different appetition responses to IG infusions of sucrose and the nonnutritive sweetener sucralose [15]. The infusions contained 16% sucrose or 1.6% sucralose, which were diluted in the gut by the consumed CS+ solution to 8% sucrose and 0.8% sucralose, respectively. These concentrations were selected because B6 mice isopreferred 8% sucrose and 0.8% sucralose in 1-min and 48-h choice tests [15,17]. Whereas the IG sucrose infusions stimulated CS+ intakes during 24-h training sessions and conditioned a robust CS+ preference, the IG sucralose infusions suppressed CS+ training intakes and conditioned a CS+ avoidance. Yet the same mice, while avoiding the CS+ flavor paired with IG sucralose, strongly preferred 0.8% sucralose to water in a 48-h two-bottle test indicating that the sweet taste of sucralose preempted the sweetener’s postoral inhibitory effects [15].

The IG infusion method provides a definitive measure of postoral sugar appetition, but a simpler, non-invasive procedure is to compare the behavioral response to orally-consumed sugars and nonnutritive sweeteners. In particular, while B6 mice isopreferred 8% sucrose and 0.8% sucralose in an initial 48-h choice Test 1 (56% sucrose preference), after separate 48-h tests with each sweetener vs. water (Tests 2 and 3), the mice displayed a strong preference (96%) for sucrose over sucralose in Test 4 [17]. This experience-induced sucrose preference is consistent with the appetition actions of IG sucrose. However, it was possible that the postoral inhibitory actions of 0.8% sucralose revealed in the IG sucralose experiment [15] contributed to experienced-induced shift in sucrose preference. We evaluated this possibility by repeating the 48-h test series with the 0.8% sucralose replaced by a sweetener mixture containing 0.1% sucralose and 0.1% saccharin (S+S). The S+S was isopreferred to 0.8% sucralose in a 1-min choice test, but unlike 0.8% sucralose, did not have postoral inhibitory actions in 48-h oral and IG infusion experiments [17]. B6 mice slightly preferred, by 60%, 8% sucrose to S+S in the initial 48-h Test 1, but after separate sugar/sweetener vs. water Tests 2 and 3, they strongly preferred (90%) sucrose to S+S in Test 4. The similar results obtained with the two different sweetener solutions indicated that the postoral inhibitory actions of 0.8% sucralose did not substantially contribute to the experience-induced preference B6 mice developed for 8% sucrose.

Quite different results, however, were obtained in sweetener vs. glucose tests. In this case, B6 mice significantly preferred 0.8% sucralose and 0.1% S+S to 8% glucose in 1-min choice tests by similar amounts (80%, 75%) [17]. Yet, in the 48-h Test 1, the mice significantly preferred glucose to 0.8% sucralose, with their preference increasing from the first to second day of the test (66 to 93%, Fig. 1A). In marked contrast, in the 48-h Test 1 with glucose vs. S+S, B6 did not prefer glucose on either test day (Fig. 1B). Yet, after Tests 2 and 3 with glucose and S+S vs. water, the mice displayed an 88% preference for glucose over S+S in Test 4, confirming the postoral appetition actions of the sugar [17]. Taken together, the findings that B6 mice preferred glucose to 0.8% sucralose but not to 0.1% S+S in the first 48-h test suggest that the postoral inhibitory actions of 0.8% sucralose contributed to the significant glucose preference observed in this test. In marked contrast to the glucose results, B6 mice tested with 8% fructose, which has no postoral appetition actions in this strain [14,22], significantly preferred both 0.8% sucralose and 0.1% S+S to fructose, by 76% and 81% respectively, in Test l [17]. Yet, sweetener preferences diverged after the mice had separate experience with fructose vs. water and sweetener vs. water in Tests 2 and 3. Now, the mice tested with 0.8% sucralose failed to prefer (48%) the sweetener over fructose in Test 4, while the mice tested with 0.1% S+S maintained their strong sweetener preference (78%) over fructose. The selective loss in 0.8% sucralose preference may have resulted because the mice experienced the postoral inhibitory actions of sucralose in the sweetener vs. water test. Given the differential results obtained with sucralose and S+S in these 48-h tests, Experiment 1 investigated if the two sweetener solutions promote differential glucose appetition using a 1-h test paradigm which is a very sensitive measure of glucose appetition [21].

**Fig. 1.**
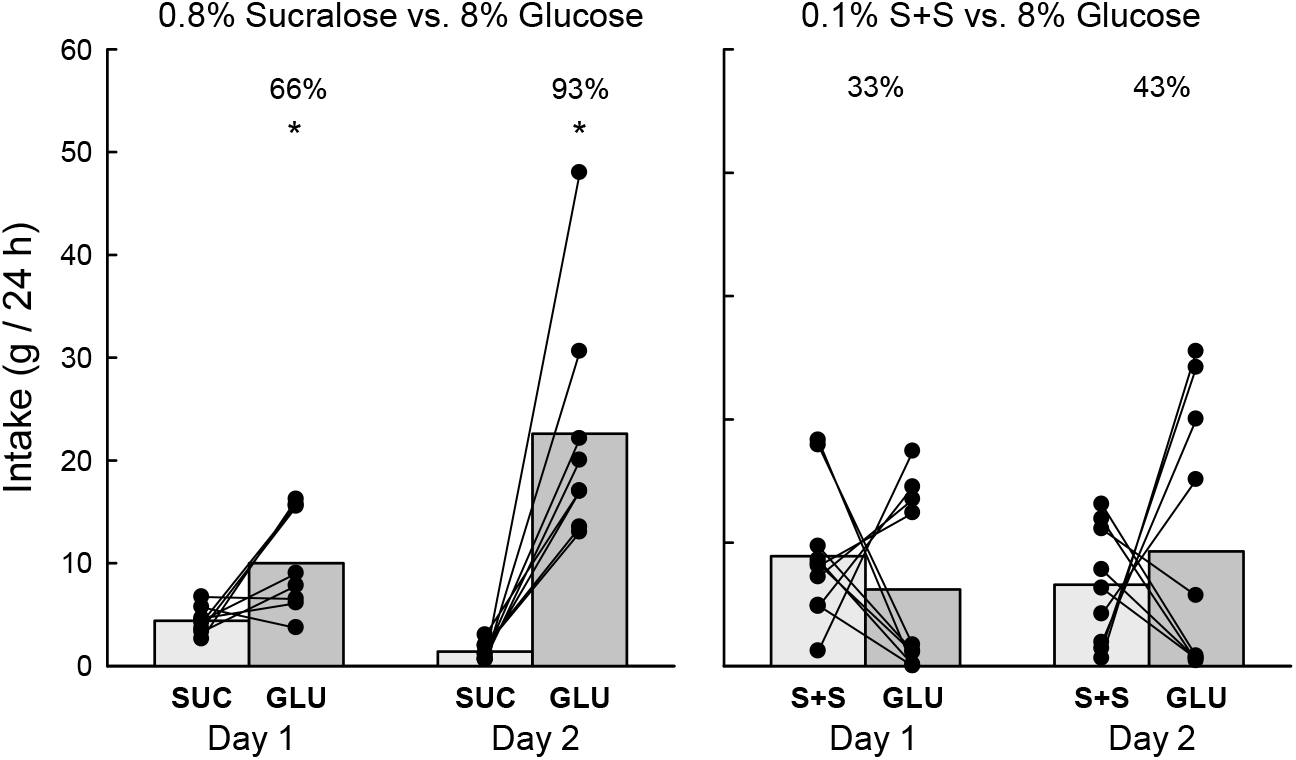
A. Group mean (bars) and individual subject (lines) 24-h intakes of 0.8% sucralose (SUC) vs. 8% glucose (GLU) during days 1 and 2 of two-bottle, 48-h Test 1 (see Figure 2 in [17]). B. Group mean (bars) and individual subject (lines) 24-h intakes of 0.1% sucralose + saccharin (S+S) vs. 8% glucose (GLU) during days 1 and 2 of two-bottle, 48-h Test 1 (see Figure 3 in [17]). Numbers atop bars represent the mean percent preference for 8% glucose. * Significant (p < 0.05) differences between SUC vs. GLU intakes.

## 2. Experiment 1. Glucose appetition with 0.8% Sucralose or 0.1% S+S sweetener solutions

In 48-h tests, ad libitum fed mice given concurrent access to nutritive and nonnutritive solutions take frequent, small drinks and consequently sugar appetition occurs slowly over time (see [3,7,18]). With food-restricted animals given one-bottle access to sweeteners in 1-h sessions, however, sugar appetition can occur within minutes. In particular, food-restricted B6 mice trained to drink a CS-flavored 0.8% sucralose solution for three daily sessions and then switched to a CS+ flavored 8% glucose solution increased their licking rate 4-6 minutes into the first CS+/glucose session and increased it further in subsequent sessions [21]. In a two-bottle choice with both flavors presented in 0.8% sucralose, the mice preferred the CS+ to CS-by 80%. These results revealed that ingested glucose rapidly generates appetition signals that stimulate ingestion and reinforce flavor preferences. Further evidence was provided in a parallel experiment in which mice were given a CS-/saccharin solution paired with IG water infusions and then switched to a CS+/saccharin solution paired with IG 16% glucose, which was diluted to 8% by the consumed CS+ solution [21]. In the first CS+/IG glucose 1-h test the mice increased their rate of licking within 15 min and even more quickly in subsequent CS+/IG glucose sessions. They then displayed a 72% CS+ preference in a two-bottle test with the CS+ and CS-saccharin solutions without IG infusions. The more rapid increased licking response observed with oral glucose than with IG glucose suggests that oral glucose sensing may facilitate glucose appetition, although other explanations were considered [21].

The findings obtained with mice given 48-h tests with 8% glucose vs. 0.8% sucralose or 0.1% S+S suggest another possibility [17]. That is, mice given daily 1-h drinking sessions with 0.8% sucralose may have limited their intake (and drinking rate) due to the postoral inhibitory actions of the 0.8% sucralose. If so, the rapidly increased licking rate when switched to glucose may result from the postoral stimulating actions of the sugar combined with the absence of the postoral inhibitory actions of the sucralose. Clearly, glucose appetition is the primary source of the increased intake displayed by mice switched from 0.8% sucralose to 8% glucose because other mice switched to 8% fructose, which has no appetition effect in B6 mice, showed an immediate and sustained decline in licking rate [21].

The present experiment determined if the glucose appetition response displayed by food-restricted mice in 1-h tests differs if they are trained to drink 0.8% sucralose, which has postoral inhibitory actions in 48-h tests, or 0.1% S+S which does not have postoral inhibitory actions in B6 mice [17].

### 2.1. Methods

#### 2.1.1. Animals

Adult male C57BL/6J (B6, n=20, 9 weeks old) mice born in the laboratory from Jackson Lab (Bar Harbor, ME) stock were singly housed in plastic tub cages in a room maintained at 22°C with a 12:12-h light-dark cycle. The mice were initially given ad libitum access to chow (5001, PMI Nutrition International, Brentwood, MO) and water. When food restricted, they were fed fixed-size chow pellets (0.5 or 1 g, F0171, F0173; Bio-Serv, Frenchtown, NJ) that allowed for precise adjustment of daily food rations. All tests were conducted 3 to 5 h after lights on (8 am). Experimental protocols were approved by the Institutional Animal Care and Use Committee at Brooklyn College and were performed in accordance with the National Institutes of Health Guidelines for the Care and Use of Laboratory Animals.

#### 2.1.2. Apparatus

Drinking tests were conducted in plastic test cages with one or two stainless-steel sipper spouts as previously described [21]. Motorized bottle holders positioned the sipper spouts in front of the cages at the start of a session and retracted them at the end of the session. Licking behavior was monitored with electronic lickometers (ENV-250B; Med Associates) interfaced to a computer. Intakes in 1-min tests were not measured because the low intakes precluded accurate measurements. Fluid intakes in 1-h tests were measured to the nearest 0.1 g by weighing the drinking bottles on an electronic balance interfaced to a laptop computer.

#### 2.1.3. Test Solutions

The nonnutritive sweetener solution contained 0.8% sucralose (SUC, Tate & Lyle; Dayton, OH) or 0.1% sucralose and 0.1% sodium saccharin (S+S, Sigma Aldrich, St. Louis, MO); the nutritive solution contained 8% glucose (GLU, food grade, Honeyville Food Products, Rancho Cucamonga, CA), and all solutions were prepared daily with deionized water. The solutions were flavored with 0.01% ethyl acetate or 0.01% propyl acetate (Sigma) as in our prior study [21]. Half of the mice had ethyl acetate added to the sucralose or S+S solution (the CS-/SUC or CS-/S+S) and propyl acetate added to the glucose solution (the CS+/GLU); the flavors were reversed for the remaining mice in each group. During flavor choice tests, both flavors were added to SUC or S+S solutions which are referred to as CS+ and CS-.

#### 2.1.4. Procedure

The mice were divided into SUC and S+S groups (n=10 each) equated for body weight. They were adapted to the test cages overnight with ad libitum access to food and water. They were then given restricted access to water (1 h/day) in their home cages and trained to drink water from two bottles in the test cages for 5 min. The next day they were given 1-min choice tests with unflavored 0.8% sucralose vs. 8% glucose (SUC group) or 0.1% S+S vs. 8% glucose (S+S group) while still water deprived. The mice were then given ad libitum access to water but a restricted food ration (1–2.5 g/day) that maintained their body weights at 85–90% of free-feeding levels. On the following 2 test days, they were given 1-min tests with sweetener vs. glucose. In these tests, the mice were first given 5-s access to one sipper tube and then 5-s access to the other sipper tube to allow them to sample both solutions before being presented with both solutions for 1 min. The timing of each session for each mouse began with its 10th lick and the bottles were automatically retracted 5 s or 1 min later. The mice were returned to their home cages after the 1-min test. One hour later, they were given a second 1-min test with the left–right position of the solutions reversed. Food rations were placed in the home cages 1 h after the last test.

The mice were then given the flavored solutions in a series of one-bottle sessions (1 h/day) with the CS-/SUC or CS-/S+S on days 1-3 and the CS+/GLU on days 4-6. This was followed by four additional 1-h sessions with the CS-/SUC or CS-/S+S (days 7 and 9) and CS+/GLU (days 8 and 10). Two-bottle choice sessions (1 h/day) were conducted on days 11 and 12 with the CS+ vs. CS-flavors both presented in SUC or S+S solutions. The left-right positions of the CS+ and CS-solutions were counterbalanced throughout testing. This test protocol was based on our prior studies [16,21]. Food rations were provided in the home cages 1 h after the sessions.

#### 2.1.5. Data analysis

Sweetener (SUC or S+S) and glucose licks during the food deprived 1-min choice tests were averaged and analyzed using a mixed model ANOVA with a group factor (SUC or S+S) and repeated-measure factor (S+S vs. glucose). CS-/Sucralose or CS-/S+S licks and intakes during 1-h tests 2 and 3 were averaged (referred to as CS-Test 0) and were compared to the licks and intakes recorded in the next three CS+/GLU sessions (CS+ Tests 1-3). The 1-h total lick and intake data were analyzed in separate ANOVAs with a group factor (SUC vs. S+S) and repeated measure factor (Tests 0-3). Sweetener/sugar licks and intakes during tests 7-10 were averaged and analyzed in separate ANOVAs as were the licks in the two-bottle test. Cumulative 60-min lick curves were generated for Tests 0 to 3, and licking rates were also expressed as licks per 3-min bin for these tests. The data from the first 10 3-min bins of Tests 0 to 3, which contained the majority of licks, were analyzed separately for each group with repeated measure ANOVA (Tests x Bins) and with each CS+ glucose test compared to sweetener Test 0.

### 2.2. Results

The SUC and S+S groups licked significantly more sweetener than glucose in the food deprived 1-min tests (F(1,16) = 99.5, P < 0.001) and they did not differ in their 1-min licks or sweetener preferences over glucose (77% vs. 80%) (Fig. 2). In 1-h Test CS-0, the two groups licked similar amounts for the CS-/SUC and CS-/S+S solutions and consumed identical amounts of sweeteners (2.1 g/h). When switched from the CS-/sweetener solution to the CS+/glucose solution the two groups increased their 1-h licks to the same degree (Fig. 2). The ANOVA revealed that both groups increased their 1-h licks from Tests 0 to 1, and then from Test 1 to Tests 2 and 3 (F(3,54) = 28.5, P < 0.001); there were no significant group or group x test differences. Analysis of the solution intake data revealed a similar pattern. The SUC and S+S mice increased their solution intakes from Tests 0 to 1, and then further increased their intakes in Tests 2 and 3 (SUC group: Tests 0 to 3: 2.1, 2.6, 3.1, 3.4 g/1 h; S+S group: Tests 0 to 3: 2.1, 2.6, 3.2, 3.5 g/1 h) (F(3,54) = 39.9, P < 0.001). The two groups also licked more CS+/glucose than CS-/SUC and CS-/S+S in the one-bottle sessions that occurred after Test 3 [F(1,18) = 40.1, P < 0.001, SUC group, 3286.2 vs. 2367.0 licks/h; S+S group, 3790.8 vs. 2702.3 licks/h). In the two-bottle choice test conducted with the flavored sweetener solutions, the SUC and S+S mice licked considerably more for the CS+ than CS-solution (F(1,18) = 103.6, P < 0.001) and displayed similar CS+ preferences (79% and 84%; Figure 2).

**Fig. 2.**
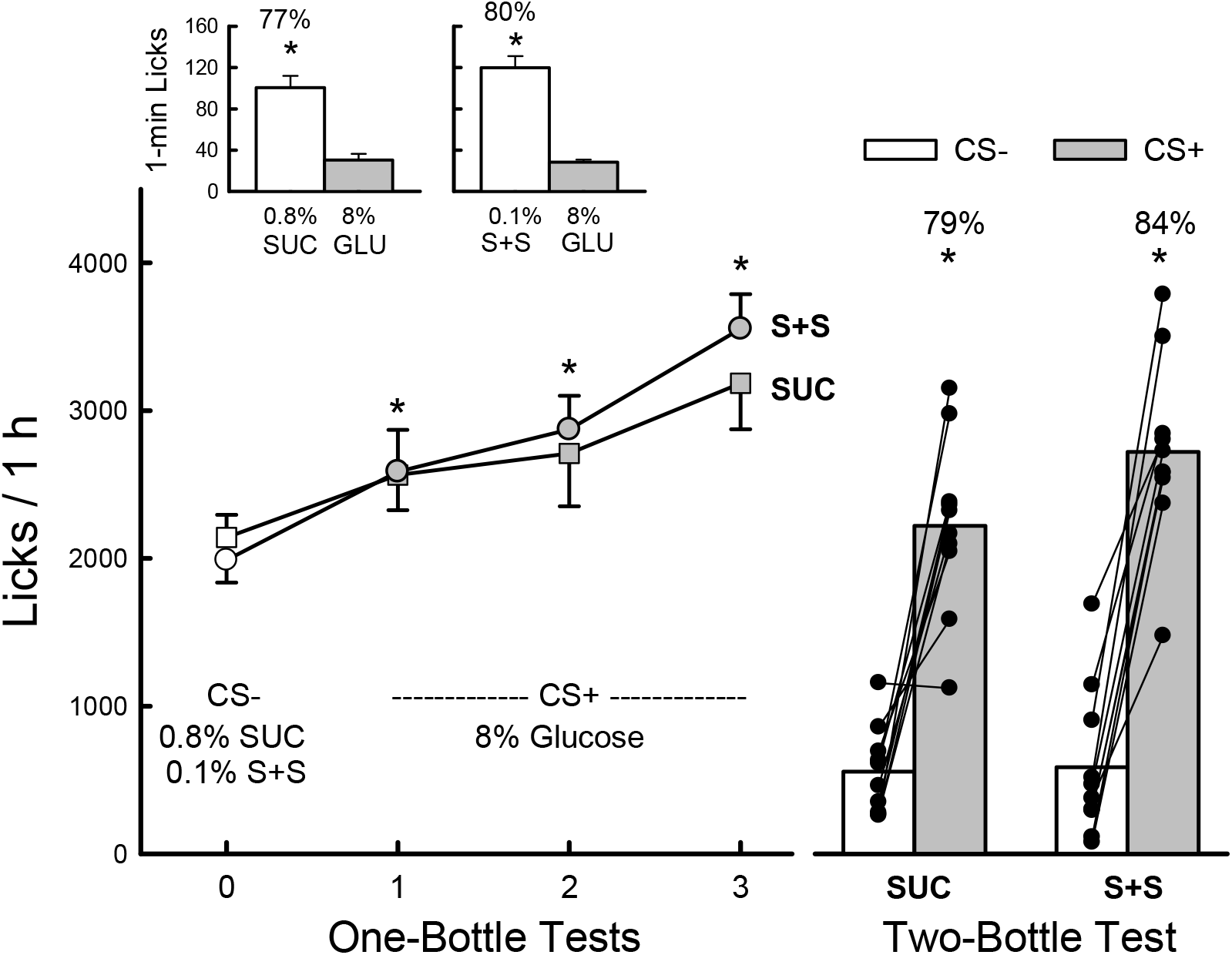
Experiment 1: Glucose stimulation of licking and flavor preferences in male mice in SUC group tested with 0.8% sucralose (SUC) and S+S group tested with 0.1% sucralose + saccharin (S+S) vs. 8% glucose (GLU). Top insets: 1-min, two-bottle choice test with unflavored SUC vs. GLU or S+S vs. GLU. Bottom left: one-bottle, 1-h licks (mean ±sem) for SUC and S+S groups tested with CS-/SUC or CS-/S+S in Test 0 and CS+/GLU in Tests 1-3. Bottom right: 1-h, two-bottle mean (bars) and individual subject (lines) licks for SUC and S+S groups tested with CS-vs. CS+ flavors presented in SUC or S+S solutions during two-bottle choice test. Numbers above bars represent mean percent preference for the specified solution. * Significant differences (P < 0.05) between SUC or S+S vs. GLU 1-min licks, Test 0 and Test 1-3 1-h, one-bottle licks, or between CS+ and CS- 1-h, two-bottle licks.

The one-bottle lick data for CS-0 and CS+1-3 tests were analyzed to determine when licking was elevated in the CS+ sessions. Fig. 3 presents the CS- and CS+ licks expressed as licks/3-min bin and as cumulative lick curves. The cumulative curves are included to show the evolution of the licking response as the animals were switched from the CS-to CS+; the statistical analysis was performed on the 3-min lick data. For the SUC mice, analysis of the bin data indicated that CS+/glucose and CS-/sucralose licks differed across tests and bins (F(27,243) = 6.7, P < 0.001). In CS+ Test 1, the mice licked less for glucose in bin 1, but more for glucose in bins 4, 5, 6, and 8 than they had for sucralose in Test 0. In Tests 2 and 3, the mice licked more for glucose than sucralose in bins 1-3 and 1-5, respectively. Similarly, the S+S group licks for glucose and S+S varied as a function of tests and bins (F(27,243) = 8.7, P < 0.001). In Test 1, the mice licked more for S+S in bin 1, but more for glucose in bins 3 to 6 while in Tests 2 and 3 they licked more for glucose in bins 1-7 and 1-8, respectively.

**Fig. 3.**
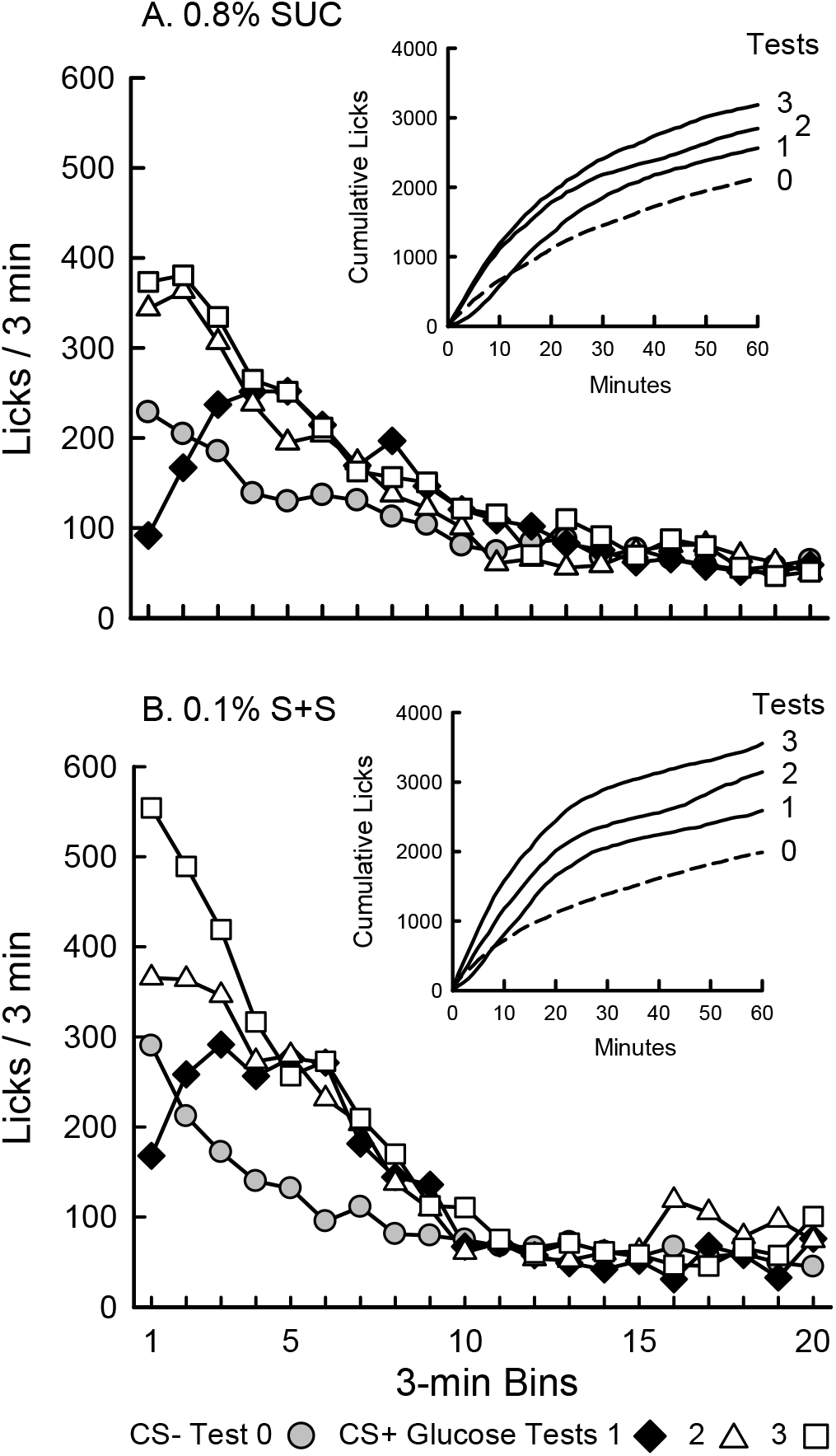
Experiment 1: Glucose-stimulated licking in SUC mice tested with 0.8% sucralose (panel A) and S+S mice tested with 0.1% sucralose + saccharin (S+S, panel B). Licks per 3-min bin are plotted for Test 0 with CS-flavored SUC or S+S and Tests 1-3 with CS+ flavored 8% glucose. Insets: graphs plot cumulative lick curves for Tests 0-3.

The present results revealed nearly identical glucose appetition responses in the mice tested with 0.8% sucralose and 0.1% S+S solutions. As explained in the General Discussion, this contrasts with the quite different results obtained with mice given 48-h tests with these solutions.

## 3. Experiment 2. Glucose appetition as a function of food deprivation state

In prior studies investigating rapid glucose-induced appetition, the mice were food restricted [16,21] but food restriction is not required to obtain sugar-conditioned preferences. In particular, in one study [2] we trained food restricted (FR) and food ad libitum (AL) B6 mice 1/h day to drink a CS-/S+S solution paired with IG water infusions and then switched them to a CS+/S+S solution paired with IG glucose infusions. Both groups displayed glucose stimulation of licking and CS+ preferences, although the effects were more pronounced in the FR mice. In the present experiment, we determined if deprivation state altered the appetition response in mice trained to drink a CS-/S+S solution and then switched to a CS+/glucose solution. One FR group and one AL group were trained with CS-/0.1%S+S and CS+/8% glucose solutions as in Experiment 1. We anticipated that the AL group would consume substantially less of these solutions than the FR group, and thus would experience less postoral glucose appetition. In order to stimulate the intakes of AL mice to approach that of the FR mice, a second AL group was trained with CS-/0.2% S+S and CS+/16% glucose solutions. A somewhat similar strategy was used in the prior IG conditioning study in that the AL and FR mice were trained with CS solutions prepared using 0.3% S+S and 0.003% S+S concentrations, respectively [2]. Female B6 mice were used in this experiment and their appetition responses were compared with those obtained with the male mice tested with 0.1% S+S in Experiment 1.

### 3.1.1. Methods

#### 3.1.2. Animals

Adult female B6 mice (9 weeks old) born in the laboratory from Jackson Lab stock were housed as in Experiment 1.

#### 3.1.3. Test Solutions

Two groups (FR8, AL8) of mice were tested with the CS-/0.1% S+S and CS+/8% glucose solutions described in the first experiment. The third group (AL16) was tested with CS-/0.2% S+S and CS+/16% glucose solutions using the flavors specified in Experiment 1.

#### 3.1.4. Procedure

The FR8 and AL8 groups (n=10 each) were adapted to the test cages overnight with ad libitum access to food and water. They were then given restricted access to water (1 h/day) in their home cages and trained to drink water from two bottles in the test cages for 5 min. The next day they were given 1-min choice tests with unflavored 0.1% S+S vs. 8% glucose while water deprived. The mice were then given ad libitum access to water but restricted food rations. On the following two test days, they were given 1-min tests with 0.1% S+S vs. 8% glucose as in the first experiment. The AL8 group was then given ad libitum access to food and water while the FR8 group was maintained on food restriction. The two groups were given two more 1-min choice tests with 0.1% S+S vs. 8% glucose in their respective food deprivation state. A third group (AL16, n=11) was tested like the AL8 group but with 0.2% S+S and 16% glucose solutions.

Following the 1-min tests, the three groups of mice were given a series of one-bottle sessions (1 h/day) under their respective deprivation state and with their respective CS- and CS+ solutions as follows: CS-/S+S on days 1-3 and CS+/GLU on days 4-6. This was followed by four additional 1-h sessions with the CS-/S+S (days 7 and 9) and CS+/GLU solutions (days 8 and 10). Two-bottle sessions (1 h/day) were then conducted on days 11 and 12 with the CS+ vs. CS-flavors presented in their respective S+S (0.1% or 0.2%) solutions. The AL8 and AL16 groups had food ad libitum in their home cages but not in the test cages. Data were analyzed as in Experiment 1.

### 3.2. Results

In the 1-min choice test while all mice were food-restricted, the three groups consumed significantly more S+S than glucose (F(2,28) = 86.6, P < 0.001); the AL16 mice licked somewhat less for the 0.2% S+S solution than the other two groups licked for 0.1% S+S but this difference was not significant (Fig. 4). In the second test conducted while the FR8 mice were food restricted, and the AL8 and AL16 mice had food ad lib, all groups again licked more for S+S than glucose (F(1,28) = 23.8, P < 0.001) and overall, the FR8 mice licked more than the AL16 mice, which in turn licked more than the AL8 mice (F(2,28) = 61.6, P < 0.001). Analysis of the percent preferences for the S+S solution revealed no significant differences between groups or the first and second tests.

**Fig. 4.**
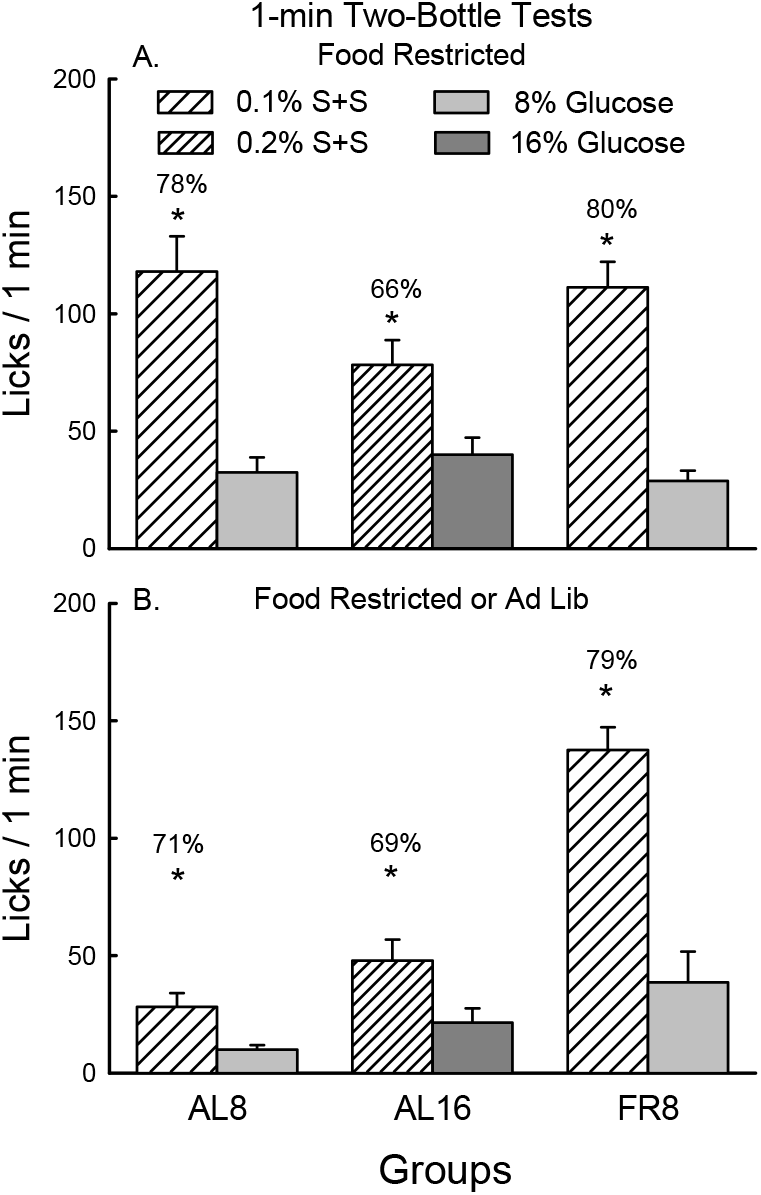
Experiment 2. A. 1-min licks in two-bottle test with FR8, AL8 and AL16 groups food restricted. B. 1-min licks in two-bottle test with FR8 group food restricted and AL8 and AL16 groups with food ad libitum. In both tests the FR8 and AL8 groups were given 0.1% S+S vs. 8% glucose while the AL16 group was given 0.2% S+S vs. 16% glucose. Numbers above bars represent mean S+S preference. * Significant differences (P < 0.05) between S+S and glucose licks.

Analysis of the CS-/S+S Test 0 and CS+/GLU Tests 1-3 revealed that, overall the FR8 group licked more than the AL8 and AL16 groups (F(2,28) =25.5, P < 0.001) and the groups increased their licks when switched from the CS-/S+S to CS+/GLU solutions (F(3,84) = 11.5, P < 0.001) (Fig. 5). However, individual tests indicated that the increases in CS+/GLU licks were significant for the FR8 and AL16 groups only. The mice also increased their solution intakes when switched from the CS-/S+S in Test 0 to the CS+/GLU in Tests 1 to 3 (F(3, 84) = 14.0, P < 0.001), but individual tests indicated that the increases were significant only for the FR8 and AL16 groups. The FR8 mice consumed more CS solutions in Tests 0 to 1-3 (1.9 to 2.2 g/h) than did the AL16 mice (1.1 to 1.6 g/h), which consumed more than the AL8 mice (0.6 to 0.9 g/h; F(2,28) = 24.6, P < 0.001). However, with respect to glucose solute intakes, overall the AL16 group consumed more glucose in the CS+/GLU tests than did the FR8 group, which consumed more than the AL8 group (0.26 > 0.17 > 0.07 g/h; F(2,28) = 45.1, P < 0.001). In the additional one-bottle test sessions that followed Test 3, overall, the FR8 mice licked more than the AL16 and AL8 mice (F(2,28) = 17.6, P < 0.001), and the FR8 group was the only group that licked significantly more for CS+/GLU than CS-/S+S (Group by CS interaction F(2,28) = 7.9, P < 0.01).

**Fig.5.**
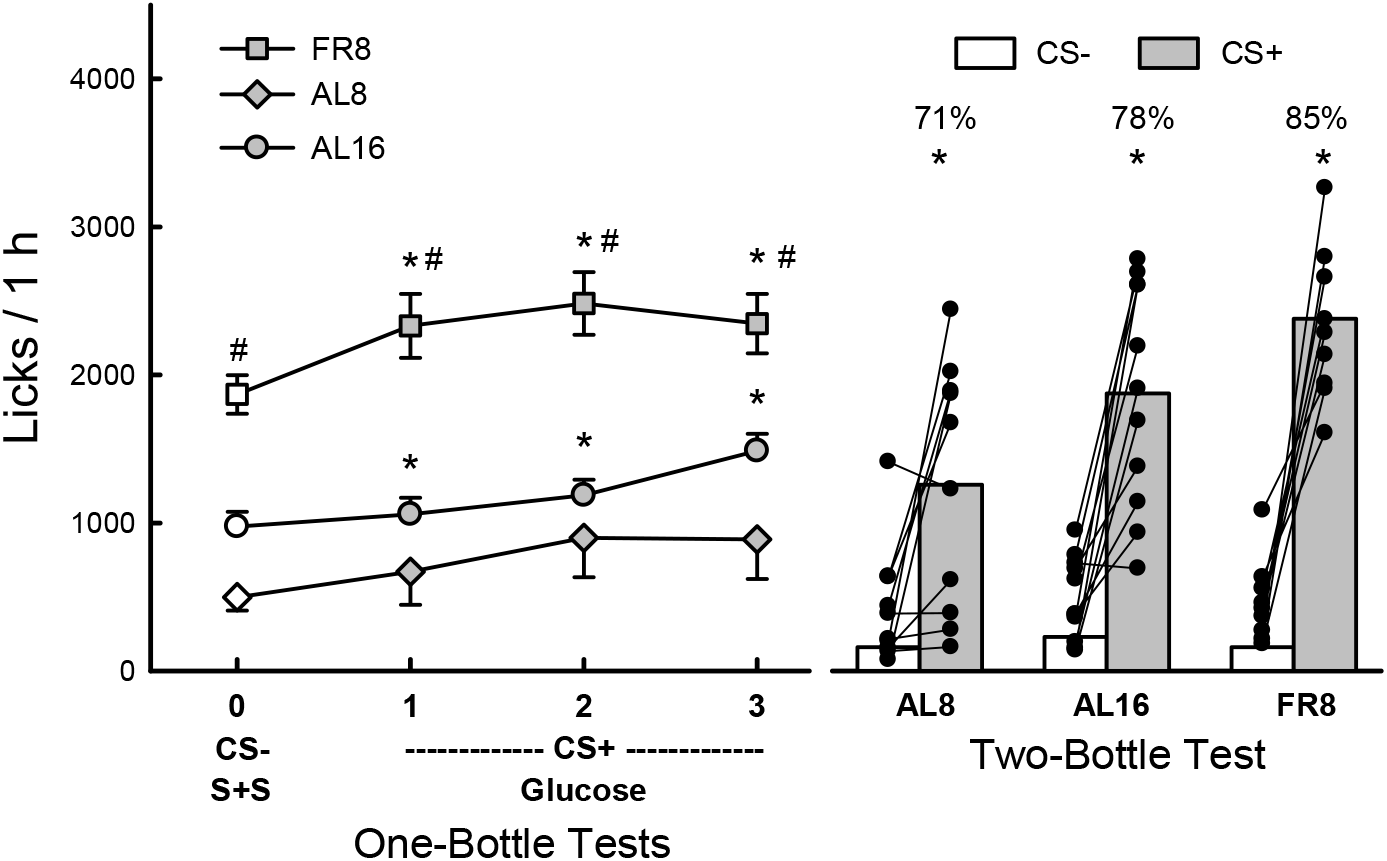
Experiment 2: Glucose stimulation of licking and flavor preferences in female FR8 and AL8 mice tested with 0.1% S+S vs. 8% glucose (GLU) and AL16 mice tested with 0.2% S+S vs. 16% glucose. FR8 mice were food restricted and AL8 and AL16 mice had food ad libitum. Left: 1-h licks (mean ±sem) for CS-/S+S in Test 0 and CS+/GLU in Tests 1-3. Significant differences (P < 0.05) between CS-/S+S licks (Test 0) vs. CS+/ GLU licks (Tests 1-3). # Significant difference between FR8 licks vs. AL8 and AL16 licks. Right: 1-h mean (bars) and individual subject (lines) licks for CS+ and CS-flavors presented in S+S solutions during two-bottle test. Numbers above bars represent mean percent preference for CS+. Significant differences (P < 0.05) between CS+ and CS- licks.

In the two-bottle preference test, all three groups licked more for CS+ than CS- (F(1,28) = 95.1, P < 0.001); also the FR8 mice licked more CS+ than did the AL8 mice (Group x CS interaction, F(2,28) = 5.0 P < 0.05) (Fig. 5). In terms of CS+ preference, the FR8 group displayed the strongest preference (85%), followed by the AL16 (78%) and AL8 (71%) groups, but these differences were not significant. A somewhat similar pattern of results was obtained for CS+ and CS-intakes. All groups consumed more CS+ than CS-(F(1,28) = 95.8, P < 0.001), the FR8 and AL16 mice did not significantly differ in CS+ intakes (1.7, 1.4 g/h), and both consumed more CS+ than the AL8 group (0.9 g/h; Group x CS interaction, F(2, 28) = 4.8, P < 0.05).

The cumulative and bin lick curves are presented in Figure 6. Analysis of the bin data for the FR8 mice indicated that they licked more for CS+/GLU than for CS-/S+S and that the difference varied across tests and bins (F(27, 243) = 4.9, P < 0.001). In Test 1, the FR8 mice licked more for CS+/GLU in bins 2 to 6 than they had for CS-/S+S in Test 0. In Tests 2 and 3, the mice licked more for CS+/GLU than CS-/S+S in bins 1-6 and 1-4, respectively. In contrast, for the AL8 mice, 3-min licks did not vary across tests but only across bins (F(9,81) =2.7, P < 0.01). The AL16 mice were like the FR8 mice in that their licks varied across tests and bins (F(27,270) = 4.3, P < 0.001). In Test 1, AL16 mice licked more CS+/GLU than CS-/S+S in bins 3, 5 and 6. In Tests 2 and 3 they licked more CS+/GLU than CS-/S+S in bins 3-5 and 1-4, respectively.

**Fig. 6.**
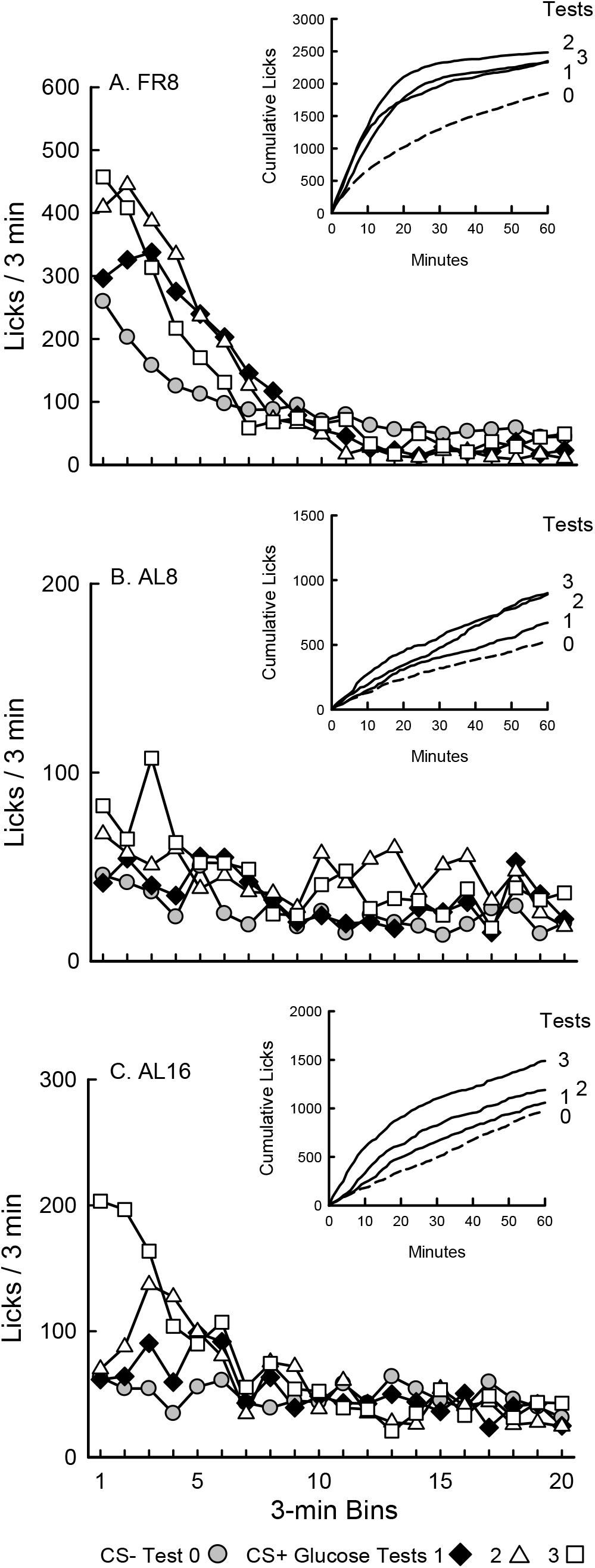
Experiment 2: Glucose-stimulated licking in food restricted FR8 mice (A) and ad libitum AL8 mice (B) tested with 0.1% S+S and 8% glucose solutions. C. Ad libitum AL16 mice tested with 0.2% S+S and 16% glucose solutions. Licks per 3-min bin are plotted for Test 0 with CS-flavored S+S and Tests 1-3 with CS+ flavored glucose solutions. Insets: cumulative lick curves for Tests 0-3.

Figure 7 compares the glucose appetition responses of the FR female mice in this experiment with the FR male S+S mice of Experiment 1. Note first that the male and female mice displayed identical preferences for S+S over glucose in the 1-min test. The two groups licked very similar amounts for the CS-/S+S in Test 0 and then increased their lick responses when given the CS+/GLU solution in Tests 1-3 (F(3,54) = 25.6, P < 0.001). They differed in that the male mice licked more than females for CS+/GLU in Tests 2 and 3 (Group x Test interaction, F(3,54) = 15.2, P < 0.001). In the two-bottle test, male and female mice were similar in that they licked more for the CS+ than CS-solutions (F(1,18) = 180.6, P < 0.001) and displayed very similar CS+ preferences (84%, 85%). Overall, the male mice consumed more sweeteners (S+S, glucose) than female mice in the one-bottle Tests 0-3 (2.8 vs. 2.2 g/h; F(1,18) = 12.8, P < 0.01) and two-bottle test (2.9 vs. 2.0 g/h; F(1,18) = 28.4, P < 0.01) consistent with the sex difference in body weight.

**Fig. 7.**
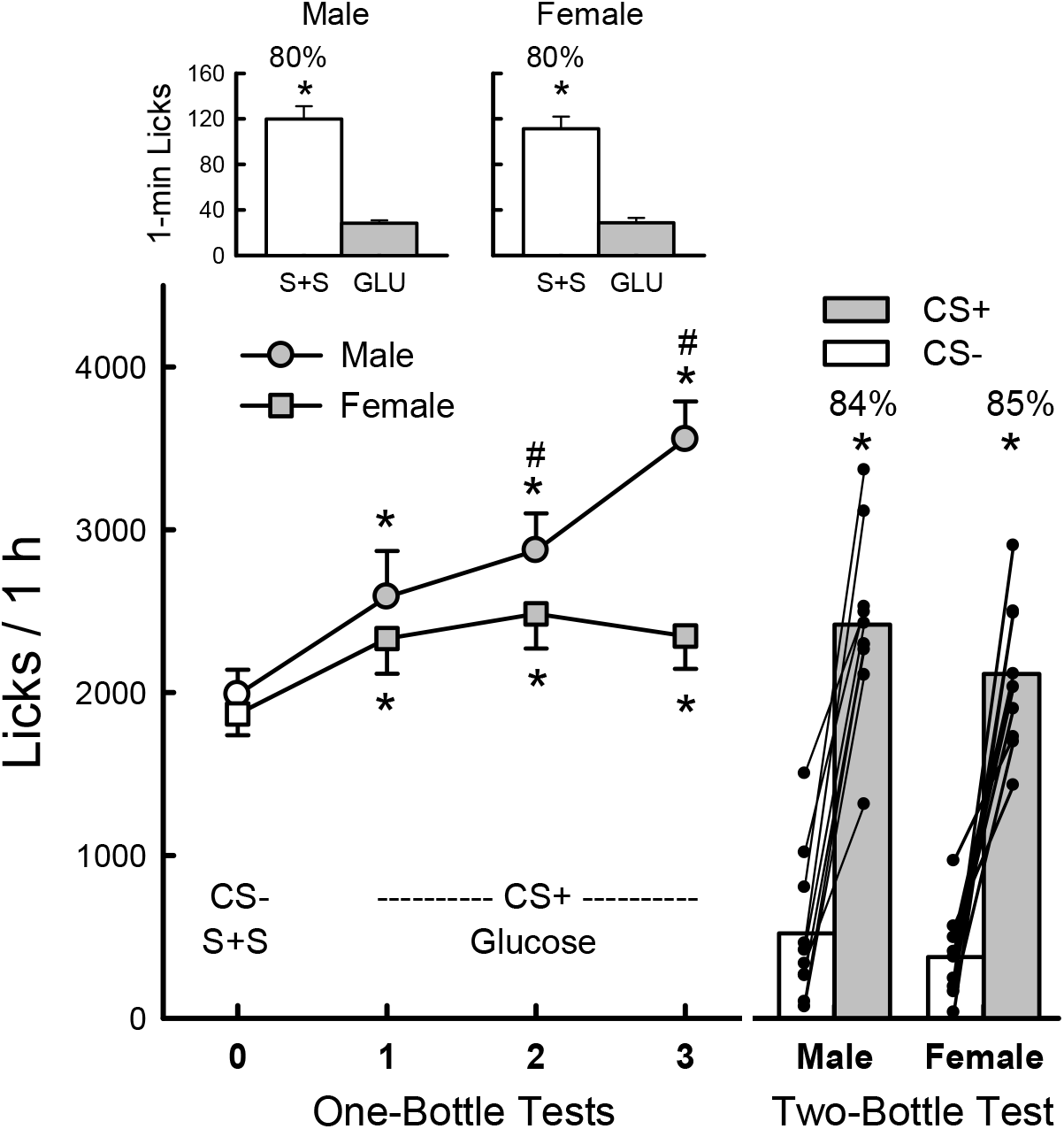
Experiments 1 and 2: Glucose stimulation of licking and flavor preferences in FR male and FR female mice tested with 0.1% sucralose + saccharin (S+S) and 8% glucose (GLU). Top insets: 1-min, two-bottle choice test with unflavored S+S vs. GLU. Bottom left: 1-h licks (mean ±sem) for CS-flavored S+S in Test 0 and CS+ flavored glucose in Tests 1-3. Right: 1-h mean (bars) and individual subject (lines) licks for CS+ and CS-flavors presented in S+S solutions in two-bottle test. Numbers above bars represent mean percent preference for specified solution. * Significant lick differences (P < 0.05) between S+S vs. GLU in 1-min test, CS-/S+S Test 0 vs. CS+/GLU in 1-h, one-bottle tests 1-3, and between CS+ vs. CS-in 1-h, two-bottle test. # Significant lick difference (P < 0.05) between male and female mice in one-bottle tests 2 and 3.

In general, the male and female mice showed similar increases in their 3-min lick responses when switched from the CS-/S+S to the CS+/glucose solutions (see Fig. 3B and 6A). One notable sex difference, however, was the initial licking response to the CS+/glucose solution in CS+ Test 1. In the first 3-min bin, the male mice, but not the female mice licked significantly less for glucose then they did for S+S in Test 0. Then the female mice licked significantly more for glucose than S+S in bins 2 to 6, while the male mice did not lick significantly more for glucose until bins 3 to 6. However, in the very first min of CS+/glucose Test 1, the females, like the males, licked less for glucose than they did for S+S in Test 0, which is consistent with the S+S preference displayed by both sexes in the 1-min, two-bottle test.

## 4. General Discussion

The present experiments investigated three potential factors influencing the rapid glucose appetition response observed in mice given 1-h tests with flavored nonnutritive sweeteners and glucose solutions: (1) the composition of the nonnutritive sweetener; (2) deprivation state, and (3) sex differences. These are discussed in turn.

### 4.1. Sweetener composition

Our original report of rapid glucose appetition in B6 mice involved training food-restricted B6 mice 1 h/day with a highly preferred 0.8% sucralose solution which was switched to a less preferred 8% glucose solution; the solutions were either unflavored or presented with CS- and CS+ flavors that allowed for flavor preference testing [21]. As replicated in Experiment 1, when switched from a CS-/0.8% sucralose to a CS+/8% glucose solution, the male mice decreased their lick rate in the first 3-min bin, which reflects the reduced palatability of glucose as revealed in the 1-min choice test. The mice then rapidly increased their licking rate in subsequent minutes of CS+/GLU Test 1. In Tests 2 and 3 they licked at even higher rates beginning in the first 3-min bin, and displayed a robust preference for the CS+ in the two-bottle test with CS flavored sucralose solutions. Although nonnutritive sweeteners are assumed to have neutral postoral reward effects, our findings that IG 1.6% sucralose (diluted to 0.8% in the gut) suppresses intake and conditions a flavor avoidance, and that adding 0.8% sucralose to 8% glucose reduces intake and preference, indicates that concentrated sucralose has postoral inhibitory actions [15,17]. This suggested the possibility that the rapid glucose appetition displayed by mice switched from 0.8% sucralose to 8% glucose may be due in part to a release from postoral sucralose inhibition. Experiment 1 revealed that mice tested with 0.1% S+S, which has little or no postoral inhibitory actions [17], displayed a glucose appetition response, i.e., rapid stimulation of licking and conditioned CS+ preference, basically identical to that displayed by the mice tested with 0.8% sucralose.

In the first experiment, the mice consumed similar amounts of 0.8% sucralose and 0.1% S+S in the 1-h CS-Test 0 (2.1 g/h) yet in our prior study mice consumed less 0.8% sucralose than 0.1% S+S in the 48-h sweetener vs. water test (5.8 vs. 11.8 g/day) [17]. Together these findings indicate that a 1-h test does not allow enough time or intake for the postoral inhibitory actions of 0.8% sucralose to be expressed. Future studies should record the one-bottle, 24-h daily drinking patterns of mice offered 0.8% sucralose or 0.1% S+S to monitor the expression of postoral sucralose inhibition.

### 4.2. Deprivation State

Prior 1-h studies [16,21] of rapid glucose appetition used food-restricted mice, but food restriction is not required to observe sugar-conditioned preferences. This is demonstrated by the sugar preferences displayed by food ad libitum mice given 48-h tests with sugars (sucrose, glucose) and nonnutritive sweeteners (sucralose, S+S, acesulfame K) [17,18]. Furthermore, IG sugar infusions condition flavor preferences in ad libitum fed mice and rats given short (0.5, 1 h) and long (24 h) sessions [2,20]. Experiment 2 specifically compared the sugar appetition responses of food-restricted and ad libitum mice given 1-h tests with flavored S+S and glucose solutions. In 1-min choice tests, the FR8 and AL8 groups strongly preferred the 0.1% S+S to 8% glucose (79%, 71%) which demonstrates the inherent palatability of the S+S solution, but the FR8 group licked much more for the sweeteners than the AL8 group, demonstrating the potency of hunger in driving sweetener intake. The FR8 mice also licked more S+S and glucose than AL8 mice in the 1-h tests, but both groups acquired significant preferences for the glucose-paired CS+ flavor (85%, 71%). The AL16 mice, which were tested with 0.2% S+S and 16% glucose solutions, showed greater glucose stimulation of licking than the AL8 mice, but their licking response was less than that of the FR8 mice. This may been due, in part, because the AL16 mice, given the more concentrated glucose solution, consumed more glucose solute than did the FR8 mice. Nevertheless, the three groups did not differ significantly in their conditioned CS+ flavor preferences. Thus, consistent with prior findings obtained in B6 mice given IG glucose infusions [2], food restriction stimulates sweetener intakes but is not required for glucose-conditioned flavor preferences.

The present findings contrast with a recent report that food deprivation is required for mice to prefer sucrose over sucralose [9]. In this study, food deprived B6 mice given a single 30-min choice test with 3.4% sucrose vs. 0.02% sucralose, which were described as being “equally sweet”, displayed a strong sucrose preference 10 min into the test session. In contrast, “well-fed” mice were reported to display no preference between these sweeteners (data were not presented). The authors proposed that the food deprived mice detected the nutritive sucrose independent of its taste and without postoral conditioning, although how they did so was not fully explained. There is accumulating evidence that mice have oral sugar sensors separate from their sweet taste receptors, but the role of these sensors in sugar preference is not well established [8,19]. An alternative explanation for the rapid preference for 3.4% sucrose over 0.02% sucralose is that the solutions were not, in fact, equally sweet. This is suggested by earlier reports that B6 mice equally preferred 4.8% sucrose and 0.4% sucralose, and 8% sucrose and 0.8% sucralose in brief choice tests [6,17]. In view of these findings, the response of food deprived and ad libitum mice to 3.4% sucrose and 0.02% sucralose solutions requires further investigation.

### 4.3. Sex differences

Most studies comparing learned sugar vs. nonnutritive sweetener preferences have used male mice. Experiments 1 and 2 investigated the glucose appetition responses of food-restricted B6 male and female mice tested with flavored 0.1% S+S and 8% glucose solutions. Overall, the male and female mice were similar in increasing their 1-h licking responses when switched from the CS-/S+S to CS+/glucose, and displayed similar conditioned CS+ preferences in the two-bottle choice test. However, the female mice showed an earlier elevated licking response to the CS+/glucose in Test 1 than did the male mice. Since the female and male mice showed identical preferences for S+S over glucose in the 1-min choice test, it is unlikely that a sex difference in sweetener taste preference was responsible for the differential early CS+ lick responses. Conceivably the female mice were more sensitive than males to the early postoral appetition signals generated by the ingested glucose, but there are many other possible explanations that need to be investigated. One recent study reported that central AgRP neuron activity has differential effects on the glucose conditioning response of male and female mice [12].

### 4.4. Neural mediation of glucose appetition and behavioral expression tests

Recent studies have revealed gut-brain neural circuits that can account for the rapid glucose appetition effects observed in this and other studies [21]. Tan et al. [18] reported that intestinal glucose infusions activated a gut-vagus-brainstem circuit that was essential for mice to learn to prefer a 10.8% glucose solution over an isopreferred 0.6% acesulfame K solution. The circuit was sugar-specific in that it was not stimulated by fructose. Consistent with prior sucralose findings with B6 mice [17,21], the mice failed to develop a preference for fructose over acesulfame K. In a subsequent study, Buchanan et al. [5] identified specific intestinal neuropod cells that rapidly activated vagal afferent fibers when exposed to intestinally infused glucose and sucrose but not fructose. Deactivation of this gut-brain circuit blocked the learned preference for 10.8% sucrose over an isopreferred 0.6% sucralose solution. Yet, while both studies demonstrated that intestinal glucose and/or sucrose rapidly activated (within seconds) the gut-vagal-brain circuit, their experiments did not provide behavioral evidence for rapid sugar appetition. In Tan et al. [18], B6 mice were given a 48-h, two-bottle test with glucose vs. acesulfame K and they did not display a strong glucose preference until after 24 h into the test. In Buchanan et al. [5], B6 mice were given daily 1-h, two-bottle tests with sucrose vs. sucralose but they did not display a strong sucrose preference until after 5 daily 1-h tests and an interposed 24-h two-bottle test. This contrasts with the glucose stimulation of licking observed 4 to 9 minutes into the first one-bottle glucose test in the present and prior studies [21]. Also earlier studies reported that rats given only one short (10- or 30-min) one-bottle test each with CS+ and CS-solutions paired with IG glucose and water, respectively, displayed a significant CS+ preference in a subsequent two-bottle test [1,11]. The delayed sugar appetition reported by Tan et al. [18] and Buchanan et al. [5] likely resulted, in part, because the mice were given two-bottle access to isopreferred sweet solutions, which makes it difficult for them to learn the postoral consequences of each sweetener. Also, the mice were not food restricted. The present and prior findings [1,11,21] indicate that one-bottle training with subsequent two-bottle choice tests is a more effective test paradigm to document rapid sugar appetition.

## Acknowledgement

This research was supported by grant DK031135 from the National Institute of Diabetes and Digestive and Kidney Diseases. The authors thank Steven Zukerman for his expert technical assistance.

